# Brain network segregation and integration during an epoch-related working memory fMRI experiment

**DOI:** 10.1101/252338

**Authors:** Peter Fransson, Björn C. Schiffler, William Hedley Thompson

## Abstract

The characterization of brain subnetwork segregation and integration has previously focused on changes that are detectable at the level of entire sessions or epochs of imaging data. In this study, we applied time-varying functional connectivity analysis together with temporal network theory to calculate point-by-point estimates in subnetwork segregation and integration during an epoch-based (2-back, 0-back, baseline) working memory fMRI experiment as well as during resting-state. This approach allowed us to follow task-related changes in subnetwork segregation and integration at a high temporal resolution. At a global level, the cognitively more taxing 2-back epochs elicited an overall stronger response of integration between subnetworks compared to the 0-back epochs. Moreover, the visual and fronto-parietal subnetworks displayed characteristic and distinct temporal profiles of segregation and integration during the 0- and 2-back epochs. During the interspersed epochs of baseline, many subnetworks, including the default mode, visual, fronto-parietal, cingulo-opercular and dorsal attention subnetworks showed pronounced increases in segregation. Using a drift diffusion model we show that the response time for the 2-back trials are correlated with integration for the fronto-parietal subnetwork and correlated with segregation for the visual subnetwork. Our results elucidate the fast-evolving events with regard to subnetwork integration and segregation that occur in an epoch-related task fMRI experiment. Our findings suggest that minute changes in subnetwork integration are of importance for task performance.

## Introduction

Research related to the brain’s large-scale functional connectome has recently been aimed to address questions pertaining to the brain’s dynamic functional connectivity [1–3]. One important aspect of the functional connectome is the balance between network segregation on one hand and integration on the other hand [4,5]. To this end, previous studies have studied fluctuations in the functional brain connectome during resting-state conditions (e.g. [6–9]). Recently, task-related changes for the temporal properties of the functional brain connectome have been targeted (for recent reviews, see [10–13]). Notably, most previous studies on the relationship between cognition and the dynamics of the functional brain connectome have either examined differences in brain connectivity when analyzed across entire epochs of data [14–16] or between separate fMRI sessions {17]. Notable is also the fact that many previous investigations of task-related changes in the functional brain connectome use versions of the sliding-window method to estimate time-varying functional connectivity [18–21]. However, sliding-window based methods have limited temporal resolution due to the necessity for analyzing large portions of consecutive data time-points (typically 30-60 seconds). Moreover, we have in a recent study shown that window-based methods fail to accurately track fluctuations in signal co-variance over time [22].

In this work we present results that reveal the temporal trajectories of brain subnetwork connectivity during 0/2-back working memory and resting-state fMRI experiments. To do this, we utilized data from the Human Connectome Project [23]. Estimates of time-varying functional connectivity were calculated per time-point using the jackknife correlation method that avoids the usage of windows and thereby provides a fine-grained temporal resolution [24]. The Power parcellation of the brain was used and the following ten subnetworks were studied (following the Cole et al., 2013 network template [25]): DMN – default mode, SM – sensorimotor, VIS – visual, FP – fronto-parietal attention, SA – saliency, CO – cingulo-opercular, AU – auditory, Sub – subcortical, DA – dorsal attention and VA - ventral attention subnetwork [26].

Our aim with the current study was two-fold. First, we tracked changes in brain network segregation and integration over time. This was quantified using a measure called Segregation Integration Difference (SID) which is defined below. Our results reveal a wide range of temporal patterns of segregation and integration between brain subnetworks during the full time-course of the working memory fMRI experiment. Moreover, we show that the degree of segregation and integration in the fronto-parietal and visual subnetworks is dependent on task complexity.

Second, we investigated the relationship between task performance and the degree of network segregation and integration. To this end, previous attempts to quantify dynamic functional connectivity to behavior include reaction times [27–30] and eye movement [30]. Regarding reaction time, dynamic measures have been correlated with the average reaction time [29] or by grouping reaction times into categories (e.g. median split, [27]) or heuristically based on task contrast [28]. In [30], a drift diffusion model was used where the model parameters were correlated with each bin of their dynamic functional connectivity estimates. This ignored the individual trial for which each dynamic network estimate originated from. A correlation between estimates of dynamic network activity with reaction time at each trial has not, to our knowledge, been shown. We found that reaction times and accuracy recorded for the 2-back trials are correlated to the dynamics of segregation and integration in the visual and fronto-parietal subnetworks.

## Results

### An illustrative example of temporal strength and Segregation Integration Difference (SID)

To describe how temporal profiles of subnetwork integration and segregation during epoch-related fMRI experiments and resting-state are quantified, we start by providing a simple example that serves to show the two central concepts used in this paper. The metrics used to quantify time-varying network properties are based on temporal network theory and an example of the usage of the two key metrics, time-evolving temporal strength and the segregation and integration difference (SID) is shown in Fig. 1. The example provided in Fig. 1A includes a temporal network that consists of two subnetworks: *A* and *B* with 3 nodes in each subnetwork and binary, undirected edges between nodes at each time-point. First, we calculate the within-subnetwork temporal strength for each subnetwork (D_A_, D_B_). This is the sum of all edge weights within each subnetwork at each time-point. Second, the value of temporal strength is then scaled by the total number of possible edges within each subnetwork (Fig. 1B). Third, we compute the between-subnetwork temporal strength (D_A,B_) as the scaled sum of edge weights that connect between nodes in the two subnetworks at each time-point (Fig. 1B). The global temporal average strength (D_global_) is added to Fig. 1B to assist visualization and it is simply the average of all edge weights for all nodes (in both subnetworks) at each time-point. To derive the degree of segregation and integration between *A* and *B* (SID_A-B_), we subtract the between-subnetwork from the within-subnetwork temporal strength (Fig. 1C). Thus, SID is constructed in such a way that a positive value implies more segregation and a negative value implies more integration between subnetworks (see Fig. 1C, in particular time-point 3 that represents a balance between integration and segregation, i.e. SID_A-B_ = 0. Time-point 7 shows a peak in subnetwork integration (i.e. the largest negative SID value). See the Material and Methods section for details regarding the jackknife correlation method, temporal strength and how to compute SID values in a general setting.

**Fig. 1.**
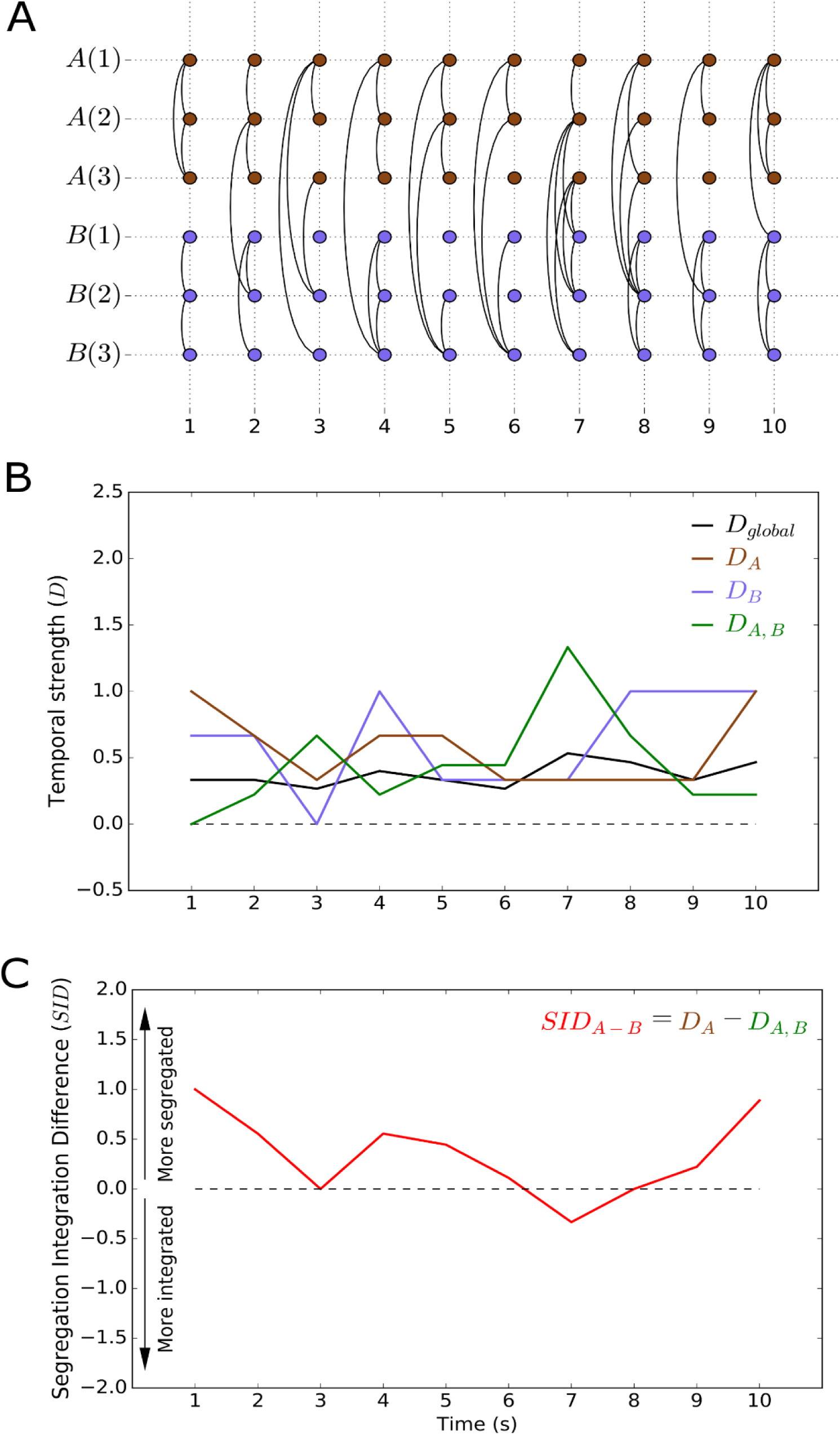
A schematic example describing the temporal network measures used. (A) A simple network example that contains two subnetworks with three nodes in each (subnetwork A and B). Connectivity (here binary) between and within subnetworks is marked with arcs as a function of time. (B) A graph that plots the value of the temporal strength at a global (D_global_) level, within-subnetworks (D_A_, D_B_) and between-subnetworks (D_A,B_). (C) The Segregation Integration Difference (SID_A-B_) measure for the two subnetworks shown in panel (A).

### Temporal strength during the working memory experiment and resting-state

The summed temporal strength over all nodes (D_Global_) during the working memory experiment and resting-state is shown for a single subject in Figs. 2A,2B and averaged across subjects in Figs. 2C and 2D. We note that D_Global_ peaks during the majority (3 out of 4) of the 15 seconds long baseline (rest) epochs in the working memory experiment while it stays constant throughout the resting-state experiment. Temporal strength at the level of individual nodes for the working memory experiment and resting-state are shown in Fig. 2E (working memory) and Fig. 2F (resting-state). Plots of temporal strength at the level of subnetworks are given in Fig. 2G (working memory) and Fig. 2H (resting-state). First, we note that the peaks in D_Global_ shown in Fig. 2C during baseline epochs for working memory experiment are not uniformly expressed across all nodes and subnetworks (Fig. 2G). Rather, while the DMN, SM, AU and VA subnetworks show peaks in temporal strength during baseline epochs, the VIS, DA, and the FP subnetworks, display the opposite pattern with a trough during baseline epochs. Second, for the resting-state experiment, small, non-uniform changes are observed for all subnetworks which suggests that the results obtained for the working memory experiment reveals time-varying connectivity that are time-locked to the experiment design.

**Fig. 2.**
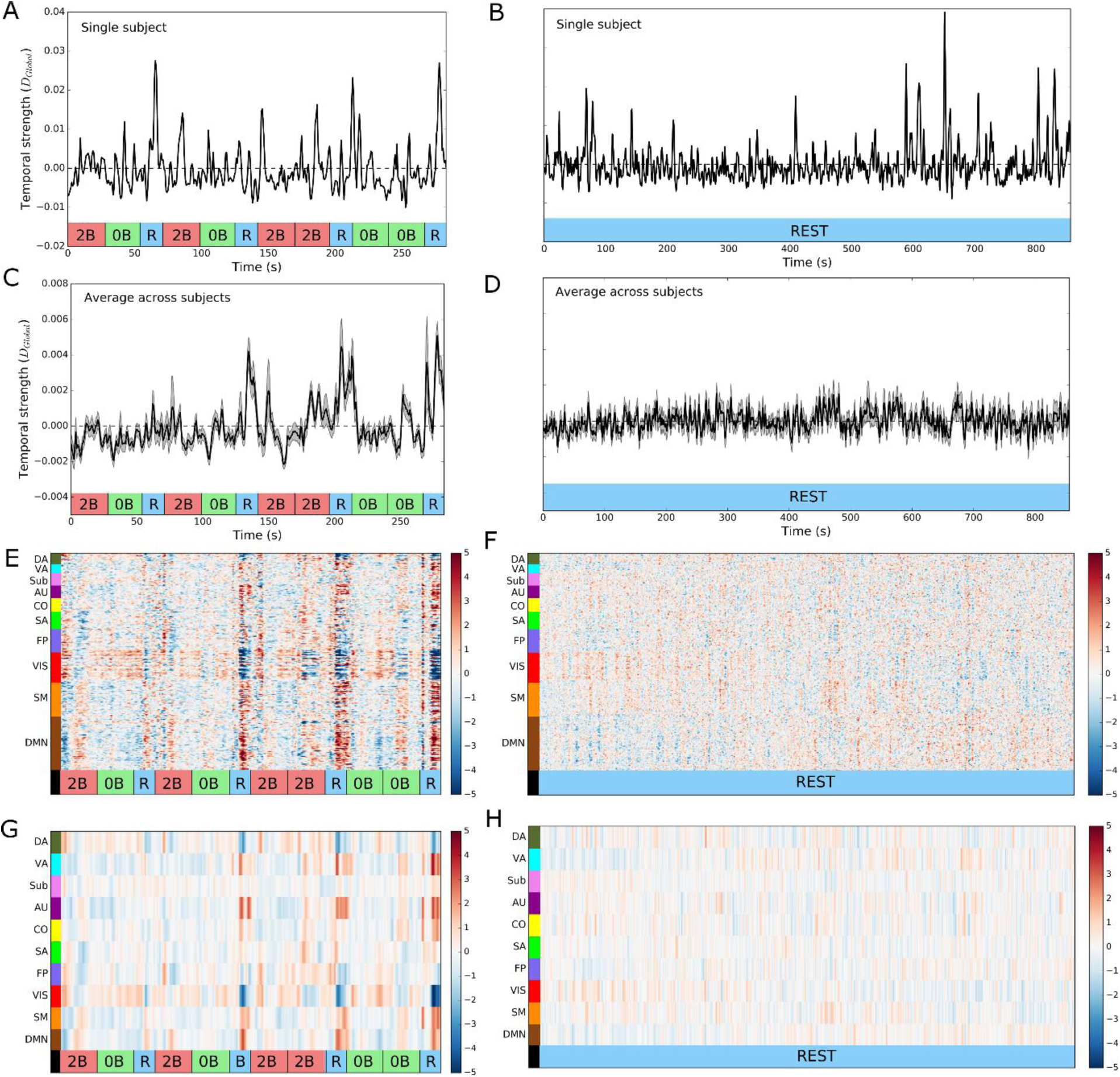
Temporal strength during the working memory experiment and resting-state experiments. (A,B) The global temporal strength (D_Global_) in a single subject during the working memory (A) and the resting-state experiments (B). (C,D) Same as in (A,B) but results are pooled (averaged across subjects) for the working memory (C) and resting-state experiments (D). (E,F) Temporal strength computed at a nodal level and averaged across subjects for the working memory (E) and resting-state experiments (F). (G,H) Temporal strength at a subnetwork level (averaged across subjects) during the working memory (G) and resting-state experiments (H). ‘0B’ – 0-back epoch, ‘2B’ – 2-back epoch, ‘R’ – rest (baseline) epoch. Shaded areas mark the standard error of the mean.

### Correlation of subnetwork temporal strength during the working memory experiment and resting-state

We wished to assess the overall degree of similarity in temporal strength over the entire imaging session (working memory and resting-state, respectively). This was done by calculating the correlation coefficient between each pair of subnetwork temporal strength time-courses in each subject and imaging session (experiment) and then taking the average across subjects. The corresponding correlation matrices are shown in Fig. 3A (working memory) and Fig. 3B (resting-state). The temporal strength time-courses for the DMN subnetwork are negatively correlated with most other subnetworks during both the working memory and resting-state experiment. Additionally, both the VIS and SM subnetworks are negatively correlated with the FP and SA subnetworks during both experiments. On the other hand, we observe strong positive correlations between subnetworks, in particular for the AU and DA subnetworks during both the working memory experiment and resting-state. Significant experiment-related differences in the time-course of temporal strength (p<0.001, Bonferroni corrected) were found between 13 pairs of subnetworks (Fig. 3C). Notably, the majority of significant differences between working memory and resting-state for fluctuations of temporal strength between subnetworks involved the DMN, VIS, SA and CO subnetworks.

**Fig. 3.**
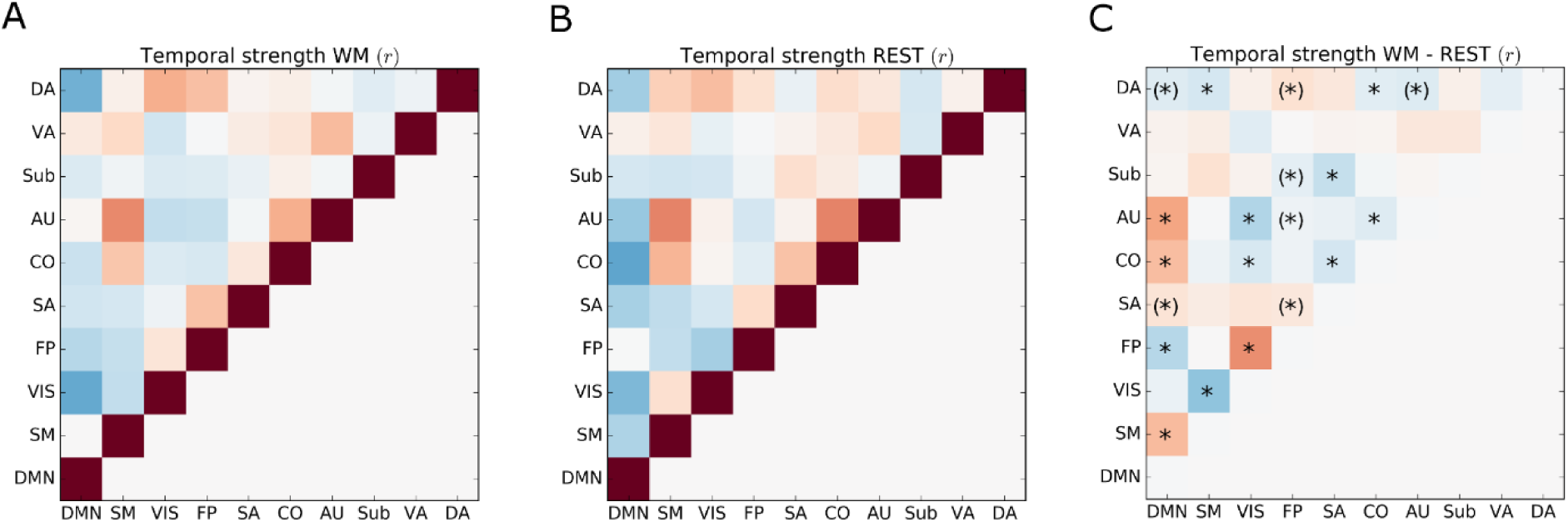
Correlation of temporal strength between subnetworks over time during the working memory and resting-state experiments. (A) Correlation coefficient matrix that show the degree of correlation in temporal strength between subnetworks during the working memory experiment. (B) Temporal strength correlation matrix for the resting-state experiment. (C) Differences in correlation of temporal strength for the working memory and resting-state experiments. Statistical significant differences in correlation were assessed by a non-parametric permutation approach (10000 permutations, p<0.001, Bonferroni corrected). An asterisk within parenthesis signify that the differences in temporal strength time-courses where only significant in one of the datasets (‘LR’ and ‘RL’, respectively).

### Subnetwork segregation and integration during the working memory and resting-state experiments

Temporal fluctuations in subnetwork integration and segregation were quantified using SID. We started by calculating SID across all subnetworks (SID_Global_, see also Materials and Methods section). Figs. 4A and 4B show the time-course of SIDGlobal in a single subject and Figs. 4C and 4D show the results obtained when averaging data across subjects. The working memory experiment time-course for SID_Global_ shown in Fig. 4C suggests a substantial global increase in segregation between subnetworks during baseline epochs. The results shown in Fig. 4C also show that the level of network integration is relatively larger during 2-back than 0-back task epochs. As shown in Fig. 4D, SID_Global_ remains rather unchanged throughout the resting-state experiment, a finding which again highlights that the observed time-varying connectivity of network integration during the working memory experiment is time-locked and tightly coupled to the epoch-related working memory experimental design.

Next, we examined fluctuations in integration and segregation at the level of subnetworks during the working memory experiment and the results are shown in Fig. 4E (single subject) and Fig. 4G (averaged across subjects). Distinct peaks in subnetwork segregation are present during baseline epochs for the VIS, DMN, FP, SA and DA subnetworks. The SID values for the VIS subnetwork show troughs during 0-back and 2-back epochs, suggesting a relative increase in integration during the working memory epochs. This is also the case, but to a lesser extent, for the DMN and DA subnetworks. Contrary to our expectations, the FP subnetwork shows a larger degree of segregation during the 2-back epochs compared to the 0-back epochs. If we consider the single subject SID results for the resting-state experiment (Fig. 4F), we observe that brief events of increased segregation occur in most subnetworks. However, those brief events of segregation are averaged out when the results are averaged across subjects (Fig. 4H). This finding is to be expected for the case when the data is not time-locked to specific stimuli or instructions.

**Fig. 4.**
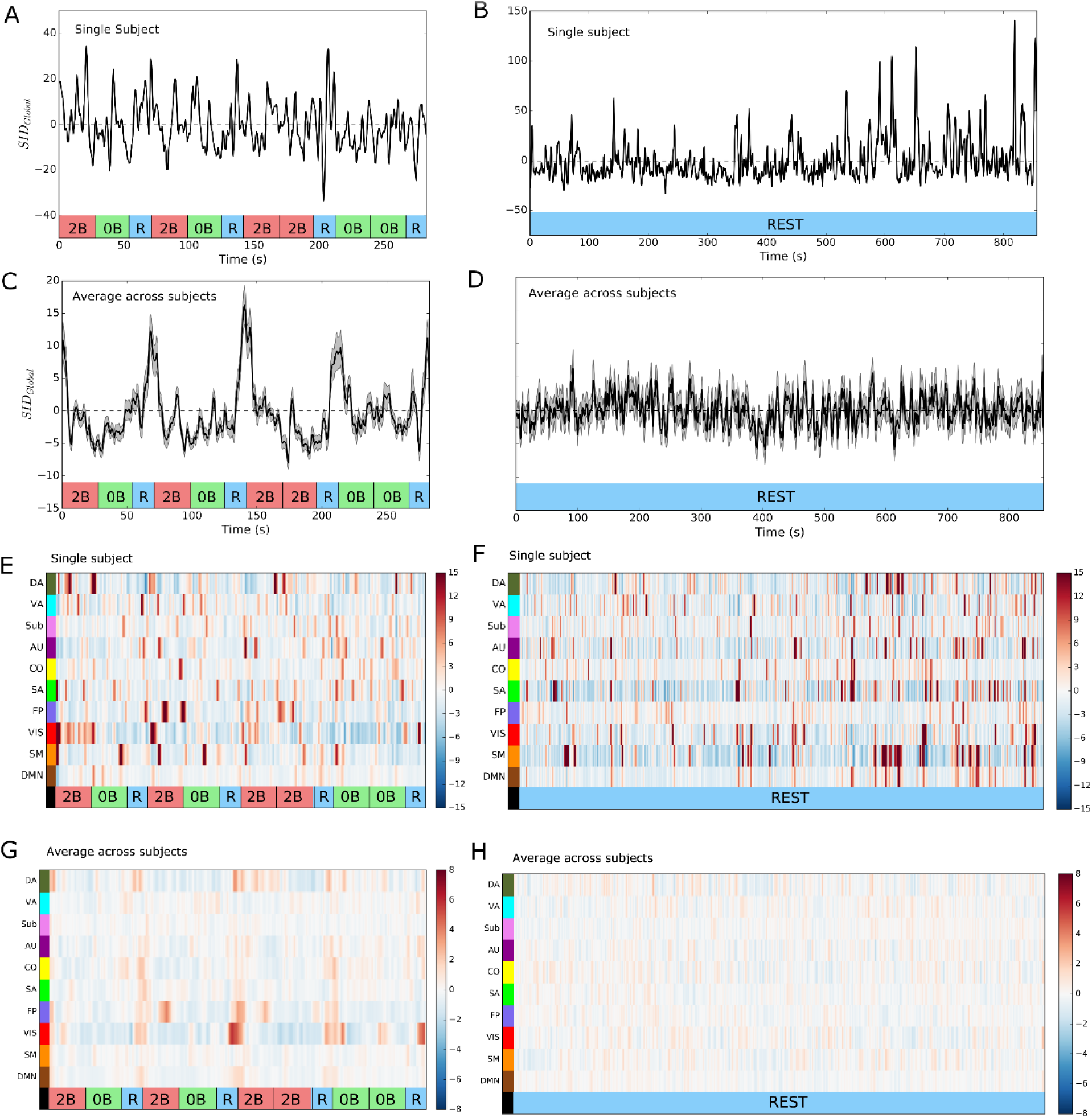
Segregation Integration Difference (SID) during the working memory and resting-state experiments. (A,B) Global SID (SIDGlobal, summed over all pairwise differences in within- versus between-subnetwork temporal strength) in a single subject during the working memory (A) and resting-state experiments (B). (C,D) Same as in (A,B) but for results that are averaged across all subjects during the working memory (C) and resting-state experiments (D). (E,F) SID results at a subnetwork level in a single subject for the working memory (E) and resting-state experiments (F). (G,H) Same as in (E,F) but for SID results that are averaged across subjects for the working memory (G) and resting-state experiments (H). ‘0B’ – 0-back epoch, ‘2B’ – 2-back epoch, ‘R’ – rest (baseline) epoch. Shaded areas mark the standard error of the mean.

### Differences in cognitive load during the working memory experiment correlates with differences in subnetwork integration and segregation

Given the difference in cognitive load and brain activity between 2-back and 0-back epochs [31–33], we were interested whether the task difficulty reflected differences in SID. This was done by dividing the working memory experiment SID time-series into 2-back, 0-back and baseline epochs. First, we examined whether there was a significant difference between the mean (first 5 seconds of data were skipped due to the hemodynamic latency) SID for the 2-back and 0-back epochs at the global level (SID_global_). The findings presented in Fig. 5A show that the mean of the time-locked and epoch-averaged SIDglobal were significantly smaller during 2-back compared to 0-back epochs (p<0.001, Bonferroni corrected). This result suggests that, at a global level, the degree of integration between subnetworks was larger during 2-back than 0-back epochs. Next, we performed the same statistical comparison for each of the ten subnetworks (Fig. 5B). Significantly lower values for the mean of SID values for the 2-back epochs compared to 0-back epochs were observed for the VIS subnetwork, suggesting a stronger degree of integration during the 2-back epoch between the VIS subnetwork and the nine other subnetworks. In contrast, SID values for the FP subnetwork were lower for the 0-back than the 2-back epochs which indicate a relatively stronger degree of integration between the FP and other subnetworks during the 0-back compared to the 2-back epochs. Additionally, the CO, SM and DA subnetworks also showed a larger degree of integration with other subnetworks during the 2-back compared to the 0-back epochs but these differences were not significant in the replication data set (see also below). The temporal profiles of the SID values for the VIS and FP subnetworks during task performance are of particular interest (Fig. 5B). Most subnetworks (DMN, SM, SA, AU, Sub, VA) are characterized by SID values that are rather constant, albeit with small fluctuations around zero, or showing a small trend of decreasing/increasing SID values (CO and DA).

### Baseline epochs in the working memory experiment is related to increases in segregation between subnetworks

The SID time-courses during baseline epochs are shown in Fig. 5C. Again, the VIS subnetwork stands out with a very strong increase in segregation that reaches a plateau after approximately 10 seconds. The DMN, FP, CO, AU and DA subnetworks shows similar increases in segregation during baseline epochs but reaches more moderate levels during the latter half of the epoch. The remaining subnetworks (SM, SA, Sub, VA) show only small or moderate fluctuations in segregation and integration during baseline epochs.

**Fig. 5.**
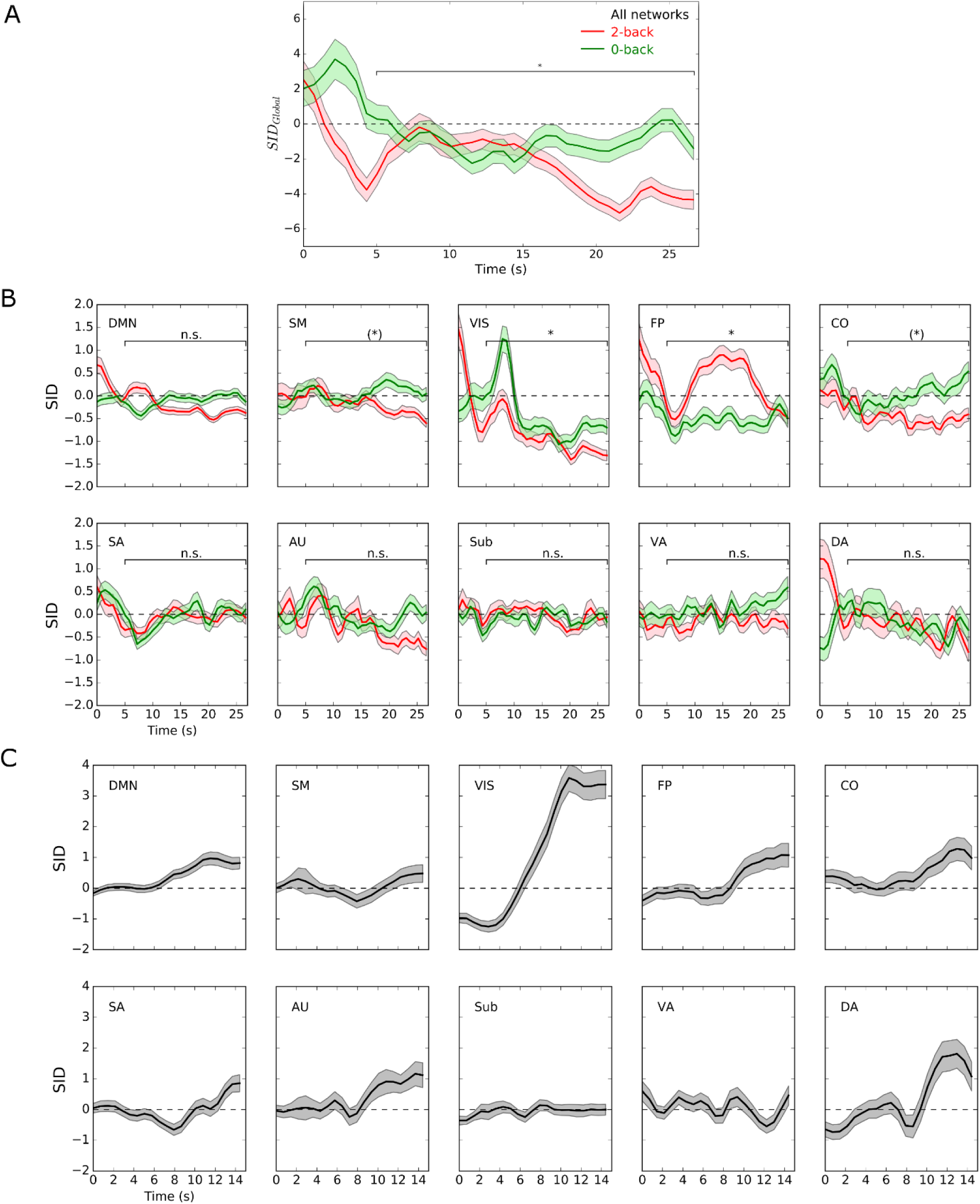
Time-locked averaging of SID results during 2-back, 0-back and baseline epochs for the working memory experiment. (A) The time-locked average (across all epochs and subjects) of SIDGlobal for 2-back (red) and 0-back (green) epochs, respectively. A non-parametric t-test (permutation approach, 10000 permutations, p<0.001, Bonferroni corrected) based on the mean of the data points from the last 20 seconds of each epoch (the first 5 seconds of data skipped due to the hemodynamic latency) was used to investigate statistically significant differences in SID during 2-back and 0-back epochs. (B) Time-locked SID results with respect to 2-back versus 0-back epochs for each of the ten subnetworks. An asterisk within parenthesis signify that the differences in SID time-courses where only significant in one of the datasets (‘LR’ and ‘RL’, respectively). (C) Time-locked averaged subnetwork SID results for the baseline epochs (results averaged across epochs and subjects). Shaded areas show the standard error of the mean.

### The degree of subnetwork integration is related to decreases in reaction time and concurrent increases in accuracy

To test the behavioral relevance of the SID results, we selected the three SID time-courses that showed consistent differences during 0- and 2-back task epochs, i.e. the SID_Global_, SIDVIS and SIDFP time-courses, respectively (see also Figs. 5A and 5B). Our aim was to investigate if the subject’s latent decision process for each 2-back trial correlated with the SID trajectories on a trial-by-trial basis. To estimate the relationship between SID results and these processes, we used a hierarchical Bayesian estimation of the drift diffusion model (HDDM) as implemented in a toolbox by Wiecki et al., 2013 (see ref. [34] and Fig. 6A for a schematic figure of the model parameters). Drift diffusion models conceptualize subject’s decisions as noisy evidence accumulation processes to fit reaction times and errors in terms of drift rate/evidence accumulation (*v*), decision boundary (*a*) and non-decision time (*T_er_*). The drift rate *v* indicates the speed of the evidence accumulation process while the decision boundary *a* relates to the response caution with which a decision is being made and the non-decision time *T_er_* refers to sensory and motor processes which are not part of the active decision process. Over the past two decades they have proved increasingly useful in psychology and neuroscience in the attempt to relate latent decision parameters to neurophysiological processes as measured for example by EEG and fMRI (see e.g. reviews found in [35,36]).

Trajectories of SID were derived by taking, for each trial, the SID value at trial onset combined with the next 9 time-points and then reduced using principal component analysis (PCA). For each of the three SID subnetwork configurations, five principal components (PC) were obtained which explained at least 95% of the variance (See Fig. 6A). The PCA weights for each of the five PCs are shown on the left side of Fig. 6B-D. A positive weight at time t is indicative of “a larger value of SID at time t” which entails more segregation and vice versa for negative weights. The PCs represent different time-locked trajectories of the SID values observed over trials. The first PC (top row in Fig. 6B-D), shows a slight peak in segregation after trial onset followed by a slight decrease. The second component starts with integration but becomes segregated with time. The third component shows an increase in segregation (except for SID_Global_ which shows the opposite pattern). The forth component has a peak in segregation followed by a peak in integration. The fifth component starts out by being integrated, then shows a peak in segregation which is followed by an increase in integration.

Due to potential problems with multi-collinearity, the SID_Global_ trajectory was run in a different HDDM model compared to the SID_VIS_ and SID_FP_ trajectories (Fig. 6A). Several models were tested in which the model parameters drift rate (*v*) and decision boundary (*a*) were allowed to co-vary, independently or together, with the SID values in the model. A selection of the best fitting model was done using the deviance information criterion (DIC). The model choice of only allowing drift rate (*v*) to vary had the lowest DIC values for the case of SID_VIS_ and SID_FP_. In the case of SID_Global_, allowing both decision boundary and drift rate to vary had the lowest DIC value, but the difference from the drift rate only model was negligible and was hence chosen for being a simpler model. In the SID_Global_ case there was no PC component where the posterior distribution correlated with evidence accumulation for (*v*) (Fig. 6B) such that 97.5% of the posterior distribution was above or below 0. For the VIS subnetwork, PC 3, where a large peak in integration occurs after trial onset, showed a significant correlation with evidence accumulation (Fig. 6C). This result entails that reaction times become faster and accuracy increases when the VIS subnetwork integrates with other subnetworks following each trial. This result makes intuitive sense as the VIS subnetwork must pass on its information regarding the visual stimuli to other brain networks for further information processing.

The FP subnetwork had two PC trajectories which had a negative correlation with the rate of evidence accumulation (PC 1 and 2) (Fig. 6D). These PC components entail that if the FP subnetwork slowly segregates throughout the trial, or if the FP subnetwork is first integrated and then segregate, this subnetwork behavior hampers task performance that leads to longer reaction times. This finding suggests that there is a sweet spot where SIDFP trajectories are optimal in the sense of delivering a quick and accurate reaction. If the FP subnetwork segregates too much it may miss important information about the trial. If the FP network integrates too much, other brain subnetworks may act as distractors. In sum, different trial-by-trial trajectories of subnetwork specific SID values for VIS and FP subnetworks could explain different aspects of the evidence accumulation parameter for the reaction time of 2-back trials in the working memory fMRI experiment.

**Fig. 6.**
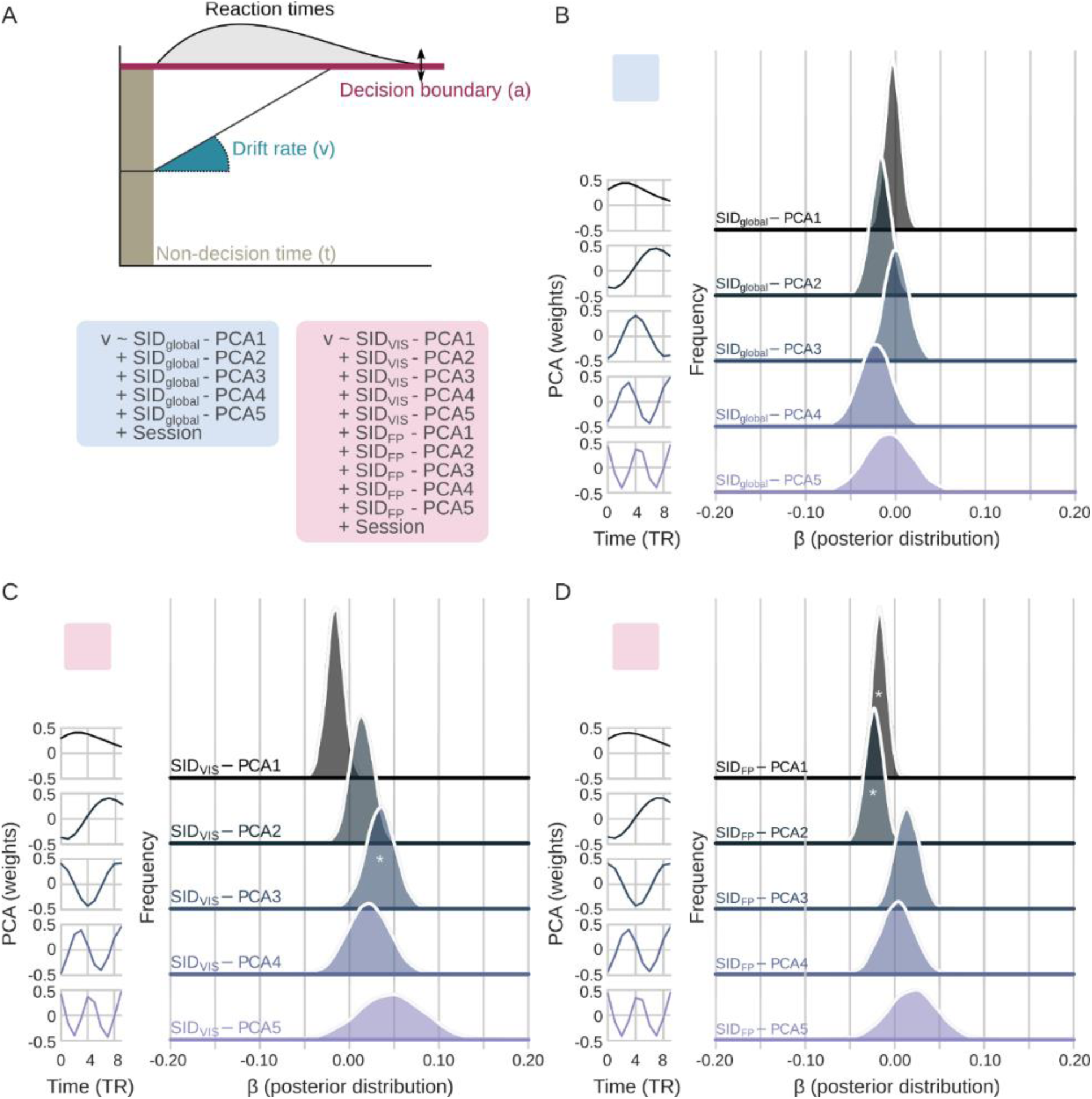
Drift rate from the Hierarchical Bayesian estimation of the drift diffusion model (HDDM) correlates with SID values related to the visual (VIS) and fronto-parietal (FP) subnetworks. (A) Schematic overview of the HDDM outlining three of the fitted parameters in the model (drift rate, decision boundary and non-decision time). Blue and red squares show the chosen two models for global SID (SID_Global_) and VIS/FP (SID_VIS_, SID_FP_) subnetwork SID. These colored squares are used in subsequent panels to indicate which model they originate from. (B) Left panel show the weights for each principal component, illustrating the trajectory of SID that comprises each component. Right panel, posterior distribution of β of the SID_global_ varying with the drift rate. (C) Same as (B) but for principal components derived from SID_VIS_. (D) Same as (B) but principal components derived from SID_FP_. For panels (B-D) a * within the distribution designates that over 97.5% of the posterior distribution is above or below 0.

### Replication of results for the ‘RL’ dataset

To study the replicability of our results, we performed an identical analysis on the ‘RL’ data-set from the Human Connectome Project [23]. Temporal strength at a global and subnetwork level for the working memory and resting-state experiments in the ‘RL’ dataset are shown in Supplementary Fig. S2 and the corresponding correlation coefficient matrices for temporal strength time-courses are shown in Supplementary Fig. S3. SID time-courses at a global and subnetwork level for the ‘RL’ dataset are shown in Supplementary Fig. S4 whereas the time-locked responses for 2-back, 0-back and baseline epochs are displayed in Supplementary Fig. S5. Supplementary Figs. S6 and S7 show the corresponding results when the jackknife method to compute dFC is replaced with the spatial distance method (LR dataset used here). Overall, our results for the ‘RL’ datasets are in good agreement with our primary analysis. Due to the fact that the performance of the HDDM model of reaction times benefits from more time-points, the RL dataset was used in that analysis together with data from the LR dataset. The effect of session was handled by including with a session covariate in the model.

## Discussion

In this study we applied a point-based time-varying functional connectivity method to fMRI data collected from 100 subjects during a working memory and a resting-state experiment. Time-varying functional connectivity time-series were analyzed using metrics taken from temporal network theory to quantify temporal strength at each point in time. This was done at three different spatial levels: individual nodes, subnetworks and at a global level. By quantifying differences in temporal strength for between- and within-subnetworks with the SID measure, we were able to derive fine-grained temporal profiles of change in integration and segregation between brain subnetworks during the working memory experiment as well as during the resting-state experiment. Several interesting properties related to how subnetworks in the brain interact with each other during task performance, baseline periods and resting-state conditions were revealed by the present findings.

### The relationship between cognitive load and segregation/integration between subnetworks

Previous research on the functional neuroanatomy for working memory have shown that large areas in the parietal, prefrontal and visual cortical areas are involved [37]. Based on the results from the current time-varying connectivity fMRI study, we suggest that increases in cognitive load are associated with a general increase in integration between subnetworks during performance of the 2-back compared to the 0-back task (Fig. 5A). Our results are corroborated by recent studies that examined the relationship between cognitive load for the n-back task and network integration using both static [15, 38-39] and time-varying functional connectivity measures [21,30,32,41]. However, as shown here, the increase in integration is not uniform across all subnetworks, which is an important novel finding. The overall result of a larger degree of integration during the 2-back epochs seems to be driven mainly by the strong degree of integration for the VIS subnetwork, but also to some lesser extent by the DA, CO and SM subnetworks. The temporal profiles of integration and segregation for the FP and VIS subnetworks during task epochs are intriguing. We suggest that the initial peak of segregation for the VIS subnetwork reflects a rather abrupt reorganization of connectivity that is initiated at the onset of the task epoch. Thus, connections between the VIS subnetwork and other subnetworks are recalibrated and reset before new connections that needs to be established for the visual processing of the succeeding trials are formed. The sustained increase in integration for the VIS subnetwork during the 0- and 2-back epochs suggests that an elevated flow of information between the VIS and other subnetworks is needed for the completion of the 0- and 2-back epochs.

The temporal profiles of segregation and integration for the FP subnetwork during 2-back epochs show almost the opposite temporal profile, including an initial trough which suggests a brief phase of integration which is followed by a strong increase in subnetwork segregation. The cognitive importance of the initial phase of integration for the FP subnetwork is presently unclear. It is known from the literature that the FP subnetwork encompasses brain areas that play key roles in cognitive processes that are central for the execution of working memory tasks such as updating of items in memory, memory maintenance of items and comparison between items kept in memory. The increased segregation of the FP subnetwork from the other subnetworks during the 2-back epochs might reflect a mechanism that tries to suppress distracting information from other subnetworks in order to optimize performance. If the FP subnetwork segregates too much from the other subnetworks however, it leads to a less efficient processing as indicated by slower reaction times and decreases in accuracy. Thus, during 2-back epochs the FP subnetwork has to find a balance where it is generally more segregated, but not too segregated, otherwise behavior in terms of reaction time will be slower and less accurate.

### Epochs of baseline during the working memory experiment introduces increases in subnetwork segregation

A striking observation in the current study is the strong overall segregation between subnetworks during epochs of baseline which are interleaved with the 0-back and 2-back epochs in the working memory experiment (Fig. 4C). From the results shown in Fig. 4C and Fig. 4G, it seems likely that the overall tendency for subnetworks to segregate from each other during baseline epochs is partly driven by the very strong increase in segregation between the VIS subnetwork and other subnetworks, but also by the marked increases in segregation for the DMN, FP, CO, AU and DA subnetworks during baseline epochs. This observation stands in stark contrast, with some notable exceptions for the VIS and FP subnetworks as noted above, to the observation that SID values that are near or below 0 during task epochs. This suggests a presence of a balanced or slightly integrative mode of communication between subnetworks during task epochs (Fig. 5B). The reason why we see the largest differences in segregation and integration for the VIS subnetwork is perhaps not overly surprising given that the working memory data analyzed here is a visually implemented fMRI experiment and we would therefore expect to see the largest temporal fluctuations integration and segregation during the time-course of the working memory experiment for the VIS subnetwork.

### Spontaneous bursts of subnetwork segregation during the resting-state experiment

The averaged results for temporal strength and SID for the resting-state experiments show no clearly defined features in time-varying brain connectivity that are consistent over subjects. Nevertheless, it is interesting to consider the profiles of temporal strength and subnetwork segregation and integration detected in single subjects as exemplified in Figs. 2B and 4B. These results suggest that during resting-state, the functional brain connectome is characterized by a moderately integrative mode of communication between subnetworks which occasionally are interrupted by episodes of network segregation. The observed fluctuations between an integrated and a segregated mode of brain function during resting-state is in line with previous time-resolved investigation of the time-varying properties of the brain connectome [6,30]. The episodes of segregation are comparatively brief in time and their appearance show similarity to the previously demonstrated burstiness of the amplitude of functional connectivity time-series in large-scale networks during rest [7]. Further work is needed to establish the idea that the interspersed events of network segregation during rest follow a long-tailed “bursty” probability distribution.

It is interesting to consider the fact that we have, as it is often done in studies of time-varying connectivity, based our time-varying connectivity analysis on a model that stipulates a static membership of nodes to specific subnetworks that do not change over time. The parcellation of the 264 nodes into 10 subnetworks used here was based on calculating gradients of connectivity profiles of resting-state brain connectivity and it should thus be pertinent for assessing dynamic functional connectivity during rest [26].

However, other studies have taken a different approach to problem of studying time-resolved fluctuations of subnetwork integration and segregation. For example, by applying time-resolved measures of network modularity to the same data as used here, it has been shown that the degree of subnetwork reorganization in the brain increases as a function of task complexity (30]. But it is conceivable that a re-organization of subnetwork membership occur during tasks. Such a re-organization of subnetwork membership can be investigated using time-varying versions of network modularity measures. The possibility to track and follow task-induced changes in network modularity at the level of single time-points represents an interesting avenue for further research. It would likely complement and extend our findings and provide further information regarding the temporal variability of brain’s large-scale networks during task performance.

In conclusion, by combining time-varying brain connectivity analysis with metrics from temporal network theory we have gained novel insights into the ongoing fluctuations in brain connectivity during an epoch-related functional neuroimaging experiment, which is an experimental design that has acted as a workhorse for functional neuroimaging research for almost 30 years (e.g. [42]). Since its inception, the inclusion of intermittent periods of rest (or other similarly low-taxing cognitive tasks) among cognitively high-taxing tasks has been an integral component in most epoch-related fMRI experimental designs due to the necessity to make comparisons of brain activity related to differences in task complexity within the same imaging experiment. While the current results are strictly speaking only valid for the working memory task investigated here, we believe that there are good reasons to assume that the marked increases in segregation between brain networks during baseline epochs observed here will generalize to most experimental designs where epochs with low or no demands on cognitive resources are interspersed with epochs of high-taxing tasks. The brain’s propensity to reduce the flow of information between networks by increasing the level of segregation during baseline epochs is compelling. The results presented suggest that during “prolonged” baseline epochs, i.e. resting-state experiments, the balance between segregation and integration among subnetworks is frequently disrupted by fast rising and falling bouts of segregation. Thus, is seems likely that during the baseline epochs, typically included in epoch-related functional neuroimaging designs, the brain’s overall subnetwork configuration returns to its default dynamical rhythm that is characterized by a never ending ebb and flood of information flow between subnetworks.

## Materials and Methods

### Data used

The 100 independent subject dataset from the Human Connectome 500 subject release was used [23,43]. For our primary analysis, we used the ‘LR’ resting state and working memory fMRI sessions. A replicability study of our results was performed on the ‘RL’ resting-state and working memory fMRI sessions. Further information regarding the MR acquisition parameters can be found in [44]. In total, the dataset consisted of 2 × 1200 (resting-state) and 2 × 405 (working memory) image volumes per subject (TR = 0.72).

### Experimental paradigms

The working memory experiment employed a version of the n-back working memory task where 2-back and 0-back epochs (each working memory epoch consisted of a 2.5 second instruction cue followed by ten trials, each with a duration of 2.5 seconds, total duration 27.5 seconds) were interleaved with epochs of baseline (duration: 15 seconds). Each fMRI session contained 8 task epochs (4 of each object type) where each epoch contained trials that consisted of visual presentations of objects of “faces”, “body parts”, “tools” and “places”. Further information regarding the details for the stimuli presentation and paradigm design is found in [45].

### fMRI data preprocessing

The fMRI data had undergone image preprocessing and removal of artifacts from non-neuronal origins by means of the FIX ICA (FMRIB’s Independent Component Analysis-based X-noisifier) data artifact rejection process which removed ICA components from the data that were considered to constitute signal contributions from white matter, cerebro-spinal fluid, head movement, cardiac and respiratory sources [46–48]. We used the Power [26] parcellation of the cortex and subcortical regions to extract BOLD signal intensity time-courses from 264 spherical Region-of-Interests (ROI, radius = 5 mm), where each ROI is assigned to a subnetwork, see also [25] for a definition of the relationship between nodes and resting-state subnetworks. The ten subnetworks studied defined are: DMN – default mode (58 nodes), SM – sensorimotor (35 nodes), VIS – visual (31 nodes), FP – fronto-parietal attention (24 nodes), SA – saliency (18 nodes), CO – cingulo-opercular (14 nodes), AU –auditory (13 nodes), Sub – subcortical (13 nodes), DA – dorsal attention (11 nodes), VA - ventral attention subnetwork (9 nodes). Thirty-eight nodes were unassigned (U). To minimize the effect of movement [49,50], image “scrubbing” was performed. The framewise displacement (FD) for each image volume was computed (rejection at FD > 0.5). Rejected volumes were deleted and missing data-point were estimated using a cubic spline interpolation [51]. On average, 1.45% of the image volumes per subject were singled out and interpolated (4 imaging sessions). Subsequently, fMRI signal intensity time-series were band-pass filtered (0.01–0.1 Hz). fMRI resting-state data from one subject contained outliers and data from this subject was not used in the analysis.

### Calculation of time-varying functional connectivity time-series

To assess temporal changes of functional connectivity we used the jackknife correlation (JC) method. The JC method was first introduced in neuroscience for the purpose of measuring single trial coherence and Granger causality on ECoG data [52] and it has recently been applied to estimate dFC in fMRI data [22]. Since we are interested in estimating BOLD signal covariance at individual data time-points, the jackknife correlation between two time-series *x* and *y* can be expressed as

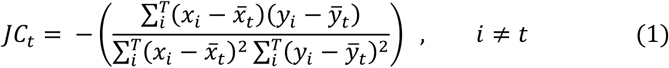

which is the Pearson correlation between *x* and *y* over all time-points with the exception of *x* and *y* at time-point *t*. The minus sign in Eq. 1 corrects for the sign inversion. The individual values of connectivity yielded by the JC method have little meaning in themselves, but, importantly, they become meaningful in relation to the rest of the time-series of dFC values. Our choice of using the JC method was based on a recent simulation study that compared five different methods to derive point-based dFC estimates [22]. In that study we showed that the JC method had superior performance in terms of tracking changes in fluctuations in signal co-variance over time [22].

Prior to computing functional connectivity time-series, we performed a linear regression on each BOLD signal time-series in which FD values and the global mean BOLD signal (taken across all ROIs at each point in time) were included as co-variates of non-interest (please also see Supplementary Text S1 for an explanation for why we regressed out the global mean BOLD signal). The FD values were included as an extra precaution against confounding effects from head-movement. The reasons for regressing out the global mean signal before computing functional connectivity time-series are discussed below. Functional connectivity time-series derived with the JC method were subsequently Fischer transformed and then converted into Z-values by subtracting the mean and dividing by the standard deviation. To assure that our results did not critically depend on our choice of method to compute functional connectivity time-series, we in parallel carried out analyses using the spatial distance method which is introduced in [24]. Further details regarding the spatial distance method is given in the Supplementary Text S2 and the results are shown in Supplementary Figures S6 and S7. Thus, the result of applying the JC method on the fMRI signal time-series was a 264×264×405 (or in the case of resting-state experiment 264×264×1200) 3D-matrix containing the time-varying functional connectivity time-series for each subject.

To apply measures from temporal network theory onto the data, we chose to treat the 264×264×405 3D-matrix as a multi-layer, temporal network where every 264×264 matrix is a connectivity matrix that represents the degree of connectivity between all nodes at time *t*. Note that the strength of connectivity between any two nodes can take any value between −1 and 1. Thus, the weights between all nodes in the connectivity matrices are undirected and weighted and take negative as well as positive values. Hence, our temporal network data model for the working memory experiment consists of 405 connectivity matrices where each individual connectivity matrix contains all available data regarding functional connectivity between all 264 nodes at each point in time. A schematic, illustrative example of a slice-plot of a temporal network representation of two simplified subnetworks, *A* (3 nodes) and *B* (3 nodes) is given in Figure 1A.

### Temporal strength

Temporal strength measures the average sum of edge weights. In the case of static networks, a node’s strength is the sum of all edge weights that are connected to it, and thus a measure of the node’s influence within the network. In the temporal case, this becomes a measure of the node’s influence within the network at time *t*. Throughout this section, equations are presented for temporal networks that contain undirected edges. We compute the temporal strength (*D*) for a node (*i*) at each time-point *t* as

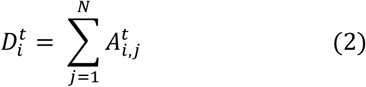

where 
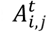
 is the connectivity matrix at time *t* and *N* is the number of nodes in the network. Temporal strength can be defined at many levels, for a single node, a subnetwork or at a global level that includes all nodes in the network. We also consider within-subnetwork and between-subnetwork temporal strength. Thus, we can at each time-point *t*, for all nodes that belong to a specific subnetwork compute the within-subnetwork temporal strength.

For example, the within-network temporal strength at time t for the subnetwork named *A* as shown in Figure 1A can be written as

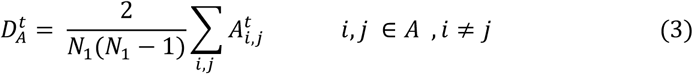

where *N_1_* is the number of nodes in *A*.

Similarly, we can estimate the temporal strength at time *t* between two subnetworks by calculating the sum of all weights associated to edges that connects between two subnetworks. For example, the between-subnetwork temporal strength for the *A* and *B* subnetworks shown in Figure 1A is

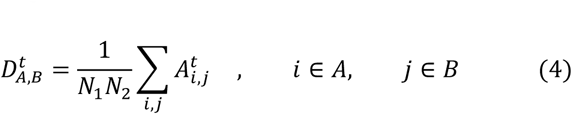

where *N_1_* and *N_2_* are the number of nodes in subnetworks *A* and *B*, respectively. Lastly, we can also define the global temporal strength at time *t* by summing the weights of edges over all nodes (*N*) as

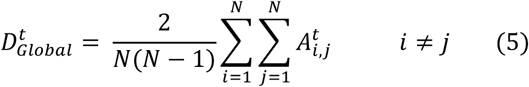

A simple and illustrative example of within-subnetwork (*D_A_*, *D_B_)*, between-subnetwork (*D_A,B_*) and global (*D_Global_*) temporal strength is given in Figure 1B.

Although all fMRI data was realigned and “scrubbed” during pre-processing, to assure that estimates of temporal strength was not contaminated by residual signal artifacts originating from head-movement, we computed the correlation between global time-series of temporal strength and time-series of FD in each subject for both working memory and resting-state experiments. The results presented in Supplementary Figure S8 show that the distribution of correlation values are centered around 0 for all four imaging sessions, suggesting that temporal strength is not biased by residual head-movement.

### Segregation and integration difference (SID)

We introduce the temporal network metric Segregation Integration Difference (SID) that operates on temporal strength at the level of subnetworks. The SID metric quantifies the degree of integration and segregation between subnetworks at each point in time. For the temporal network example shown in Figure 1, we define SID between subnetworks *A* and *B* as

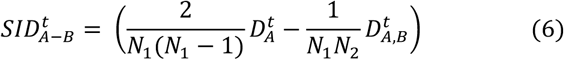

where *D_A_* and *D_B_* are calculated according to Eq. 3 (i.e. within-subnetwork temporal strength), *D_A,B_* is the between-subnetwork temporal strength as given in Eq. 4 and *N_1_* and *N_2_* are the number of nodes in the *A* and *B* subnetworks, respectively (see also Figure 1B for an illustrative example). Note that Eq. 6 is not symmetric, i.e. 
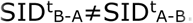
.

It is possible to extend the scope of the SID measure so that we can compute an estimate of integration-segregation between one subnetwork and all other subnetworks. If we assume that the whole brain network is divided into a total of *M* subnetworks, the SID parameter for subnetwork *m* with respect to all other subnetworks is defined as

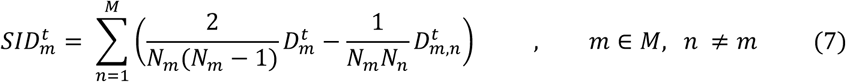

where the 
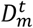
 terms are the within-subnetwork temporal strengths, the 
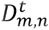
 terms represent the between-subnetwork temporal strengths and *N_m_* and *N_n_* are the number of nodes in subnetworks *m* and *n*. It then becomes possible to compute SID at a global level by summing over all subnetworks:

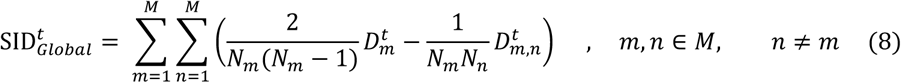

All forms of SID values defined above are derived as a pair-wise summation of differences of within-subnetwork and between-subnetwork temporal strengths (Eq. 7). Thus, an increase in SID suggests increased segregation between subnetworks that are driven either by an increased within-subnetwork temporal strength for the given subnetwork or a decreased between-subnetwork temporal strength with regard to all other subnetworks. From this follows that a decrease in SID imply an increase in integration, which is caused either by decreased within-subnetwork temporal strength and/or an increased between-subnetwork temporal strength.

The schematic example described above included only binary edges, whereas weighted edges are used in the analysis in the paper (i.e. both positive and negative values are possible for the weighted edges). It is important, for the sake of the interpretation of the results, to clarify what time-varying estimates of brain connectivity with this method entails. By definition, a normalized jackknife correlation value of 0 between two time-series at time *t* means that the correlation at *t* is the mean correlation of the entire functional connectivity time series, which approximates to the static functional connectivity between the two. In other words, whenever an estimate of time-varying connectivity is positive, it should be interpreted as time-points where the connectivity is larger than usual, and vice versa for negative values. The implication of this for temporal strength is that a negative value of temporal strength at time *t* entails that the strength of dynamic connectivity for a node at time *t* is smaller than its mean value taken across time. Thus, the usage of the jackknife method combined with the SID measure implies that a SID value of zero at time t means that the degree of segregation/integration at that point in time is approximately equivalent to its average value for the whole imaging session. When the value of SID increases, it implies more segregation than the average value across the imaging session and when it decreases it implies more integration than the average value across the imaging session.

All time-varying functional connectivity analyses (jackknife correlation, temporal strength, SID) were carried out using teneto, a package for Python ([24]; https://github.com/wiheto).

### Testing for differences in SID during 2-back and 0-back epochs in the working memory experiment

To test for significant differences in subnetwork segregation during 2-back and 0-back epochs, we averaged together SID values for different time-points within each epoch. Due to the hemodynamic latency of fMRI BOLD signals and the fact that the temporal order of task epochs was not counterbalanced with respect to baseline epochs, we did not include data from the first 5 seconds of each epoch. Statistical testing for significant differences in the time-locked and averaged (within epochs) SID values between 2-back and 0-back epochs was performed using a permutation approach (10000 permutations, p<0.001, Bonferroni corrected for multiple comparisons).

### Modeling the relationship between reaction time and SID

We sought to explore whether subnetwork SID metrics that significantly differed for the 2-back versus the 0-back epochs could explain variability in reaction times. Based on the results shown in Figures 5A and 5B, we chose the SID_VIS_, SID_FP_, SID_Global_ trajectories for further analysis. We chose here to only include the 2-back trials in our analysis (80 trials per subject), since the influence from differences in brain network segregation and integration on reaction time is expected to be larger for the more taxing 2-back trials.

As the exact time-point when SID values are relevant for behavior is uncertain, we had the choice of either reducing the number of temporal dimensions of the data, or alternatively, to run the analysis per time-point. We opted to choose the dimensional reduction route. Ten time-points were used for each 2-back trial (onset data-point plus 9 additional time-points). Principal Component Analysis (PCA) was performed for each subnetwork SID trajectory. For all three reduced datasets (SID_VIS_, SID_FP_, SID_Global_), five principle components explained 95% of the variance. Thus, five principle components could be said to accurately represent trajectories of SID time-courses after each 2-back trial. Due to the sluggishness of fMRI BOLD response, SID data at each trial onset would reflect activity prior to trial onset (and reveal possible anticipatory effects).

We employed two different models: the first contained SID trajectories from the VIS and FP subnetworks and the second model used only the SIDGlobal trajectory data. In the former case, we combined the information from both SID_VIS_ and SID_FP_, as they displayed a low correlation with each other (mean absolute r of corresponding SID_VIS_ and SID_FP_: 0.025), whereas the SID_Global_ data had to be treated separately in another model because it partly was correlated with both SID_VIS_ and SID_FP_ (mean absolute r of corresponding SID_Global_ with SID_VIS_: 0.308 and with SID_FP_: 0.309), which might cause difficulties with colinearity in the model. To this end, we created two different models where reaction time and accuracy were modelled with a hierarchical Bayesian estimation of the drift diffusion model (HDDM, [34]). The first model used ten PC components (five components from each of SID_VIS_ and SID_FP_, respectively) and the second model used five PC components extracted from SID_Global_.

The temporal trajectories of SID measures were used as regressors in the HDDM model where both the drift rate (*v*) and the decision threshold (*a*) could vary with the respective SID co-variates. The three combinations of parameters (for each covariate set) were allowed to vary in the model, (i) the decision threshold (*a*), (ii) drift rate (*v*) and (iii) both decision threshold and drift rate (*a+v*) varying with respect to the covariates. 2-back trial data from both datasets (RL and LR) were used in the same model. A binary regressor was specified to differentiate between the two fMRI sessions in both models. The reasoning for including both datasets in the same model was twofold: (1) more data makes the HDDM model more robust, (2) there are possible learning effects between the RL and LR datasets which affect both the median reaction time and accuracy, making a true replication difficult.

In the main text the results pertaining to allowing the drift rate (*v*) to co-vary with the regressors was chosen. For the VIS/FP subnetwork model, this model had the lowest Deviance Information Criterion (DIC) value (see Supplementary Table 1). In the global model (SID_Global_), the drift rate (*v*) model had a slightly larger DIC value than the combined decision threshold and drift rate (*v+a*) model. Since the difference between the two models was small and the drift rate (*v*) only is the simpler model, the drift rate model was chosen for the main text. MCMC (Markov Chain Monte Carlo) was run where 11000 samples were drawn and the first 1000 samples were burned. We used the standard priors implemented in HDDM and based on previous studies collected by [53]. Verification of chain convergence was done for all models through visual inspection and, for the chosen best fitting model for each covariate configuration, the Gelman-Rubin statistic was close to 1 for all fitted models parameters after running the model five times.

## Acknowledgements

P.F. was supported by the Swedish Research Council (grant No. 2016-03352) and the Swedish e-Science Research Center. Data were provided by the Human Connectome Project, WU-Minn Consortium (Principal Investigators: David Van Essen and Kamil Ugurbil; 1U54MH091657) funded by the 16 NIH Institutes and Centers that support the NIH Blueprint for Neuroscience Research; and by the McDonnell Center for Systems Neuroscience at Washington University. The funders had no role in study design, data collection and analysis, decision to publish, or preparation of the manuscript. The authors thank Neda Kaboodvand for helpful comments on the manuscript.

## 5 Supporting information.

### S1 Text

#### The global fMRI BOLD signal and time-varying functional connectivity

We would like to draw the reader’s attention to a methodological aspect of our analysis that is of importance. In case of resting-state fMRI, the practice of regressing out the global mean brain signal (global signal regression, GSR) before calculating functional connectivity is well documented in the literature and the advantages and disadvantages of its usage has been thoroughly discussed in the recent functional neuroimaging literature (Fox et al., 2009; Saad et al., 2012; Murphy & Fox, 2017). Regarding the usage of GSR on resting-state data, previous results suggests that the global signal most likely contains contributions from both non-neuronal and neuronal signal sources, although a recent study has argued that the large majority of the global signal stems from non-neuronal physiological processes (Power et al., 2017a,b).

At first sight, there might not be any compelling reason to perform GSR on task-based fMRI data in the case of time-varying connectivity analysis since we are extracting BOLD fMRI signal time-courses from ROIs located exclusively in gray matter. However, given the previous debate regarding its usage in resting-state functional connectivity studies and that its impact in the context of time-varying connectivity has not been previously investigated, we decided to calculate temporal strength on both GSR and non-GSR regressed fMRI time-series. The relationship between the global mean BOLD signal (averaged across all 264 ROIs and all subjects) and global temporal strength (D_Global_, calculated without GSR performed on the BOLD signal intensity time-courses) during the working memory task is shown in Supplementary Figure S1A. The time-courses displayed in Supplementary Figure S1A suggests that the onsets of 0-back as well as 2-back epochs are accompanied by a strong and brief peak in global temporal strength (marked in blue in Supplementary Figure S1A). In contrast, D_Global_ exhibits its lowest values during baseline epochs. The behavior of the global temporal strength is mirrored both at a nodal (Supplementary Figure S1B) and subnetwork level (Supplementary Figure S1C), showing synchronous increases and decreases in most nodes and subnetworks. If we compare D_Global_ with the time-course for the global mean BOLD signal (marked in red in Supplementary Figure S1A), two observations can be made. First, the majority of the peaks for D_Global_ are accompanied by similar peaks in the global mean BOLD signal. Second, for the peaks of temporal strength that are not accompanied by peaks in the global mean signal (i.e. during the baseline epochs) we find troughs of the global mean BOLD signal. The second observation can be explained by the fact that marked changes in the global mean brain signal, may it be positive or negative, implies that a large majority of the nodes are affected by the same driving signal source. The fact that a large majority of the nodes experiences the same signal increase/decrease will in turn increase their functional connectivity. Hence, a decrease in the mean global BOLD signal will be reflected by an increase in global temporal strength.

The results shown in Supplementary Figure S1 suggests that estimates of time-varying functional connectivity such as temporal strength are strongly influenced by the global BOLD signal and that the large majority of nodes show very similar connectivity profiles throughout the working memory task. Hence, it is very likely that smaller, individual changes in connectivity that occur at a nodal and subnetwork level, which might be of pivotal interest for an assessment of subnetwork integration and segregation, are disguised by the global mean BOLD signal. In the context of time-varying brain connectivity during task performance, we chose to regard the mean global signal as a nuisance variable. We agree with the recent interpretation proposed by Nalci et al., 2017 that GSR can be seen as a temporal down-weighting of the segments of the BOLD signal that have large global amplitudes. Given that temporal strength during the working memory fMRI experiment is strongly correlated with changes in the global BOLD fMRI signal as shown in Supplementary Figure S1, and that the study of the temporal properties of subnetwork integration and segregation is the focus in the current work, we present results for which the GSR step has been included in the data pre-processing pipeline.

### S2 Text

#### The Spatial Distance method to compute time-varying functional connectivity time-series

To assure that our findings were not biased by a particular choice of method to compute point-based estimates of functional connectivity, we performed all connectivity estimates and network integration and segregation analyses using a different method (the spatial distance method, SD). The SD approach estimates time-varying connectivity by calculating a weight vector for each time-point based on the distance between spatial components (i.e. nodes). It computes the weights (*w*) for each time-point (*t*) by taking the inverse of each ROI’s Euclidean distance (*D*) to all other time-points (*u*) as

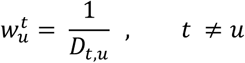

Thus, each data time-point is associated with a unique weight-vector that has a length that equals the total length of the data time-series. The weight-vectors at each time-point are subsequently scaled between 0 and 1 and the “self-weights” 
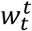
 are set to 1. The connectivity estimate at time-point (*t*) is the weighted Pearson correlation where each time-point is weighted by *w*^*t*^. The complimentary results obtained by the SD method are shown in Supplementary Figures S6 and S7. Further examples and details of the use of the SD method to estimate time-varying functional connectivity is found in Thompson et al., 2017a.

## Supplementary Figures

**Fig. S1.**
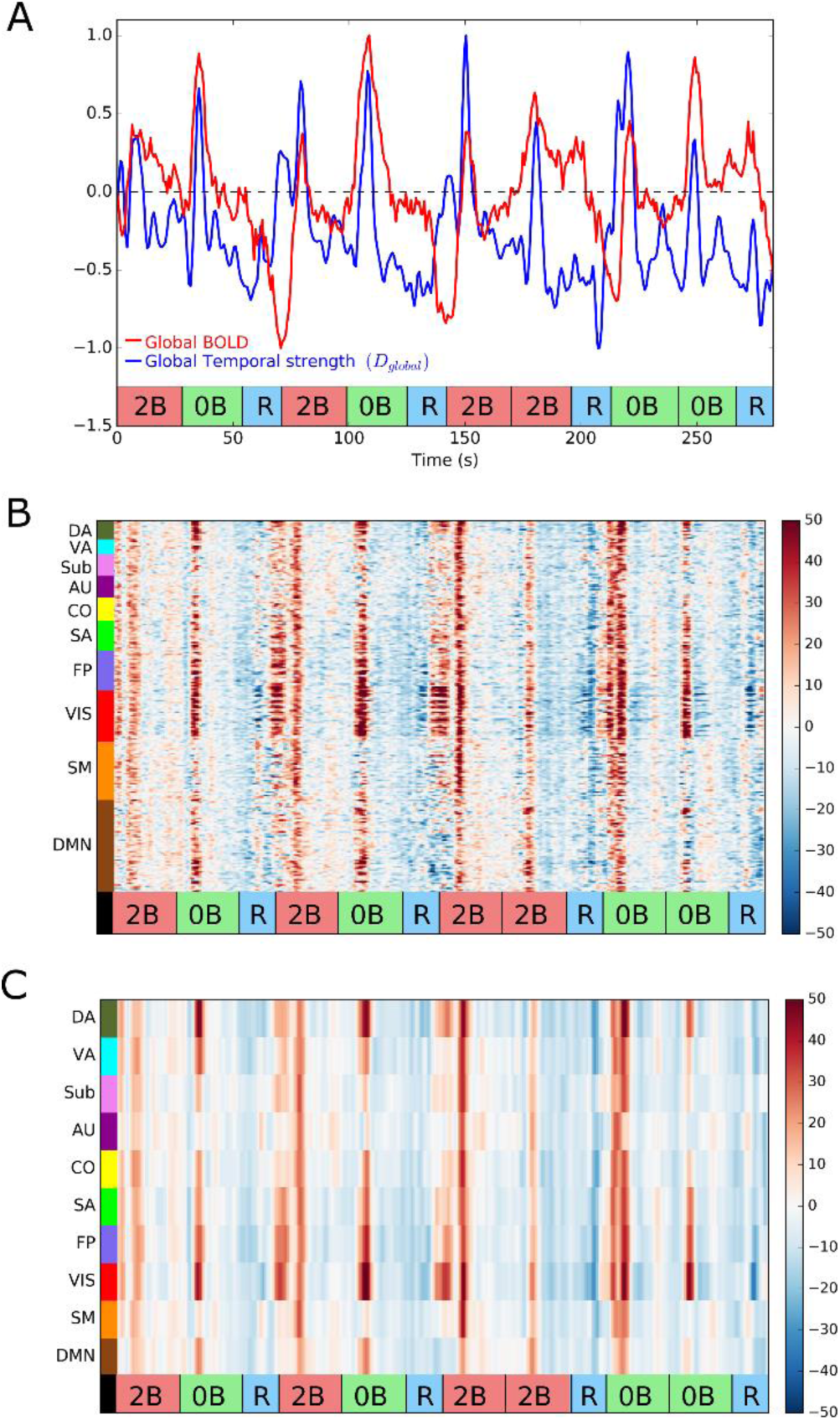
The relationship between the mean global BOLD signal and temporal strength. (A) The mean BOLD signal (averaged across all 264 ROIs and all subjects) and the global temporal strength (averaged across all subjects) for the 2-back/0-back/baseline working memory experiment. (B) Fluctuations in temporal strength at a nodal level during the working memory fMRI experiment. (C) Fluctuations at a subnetwork level during the working memory fMRI experiment. Note that the mean global BOLD signal was not regressed out prior to the computation of dynamic functional connectivity time-series and temporal centrality shown in panels (B) and (C).

**Fig. S2.**
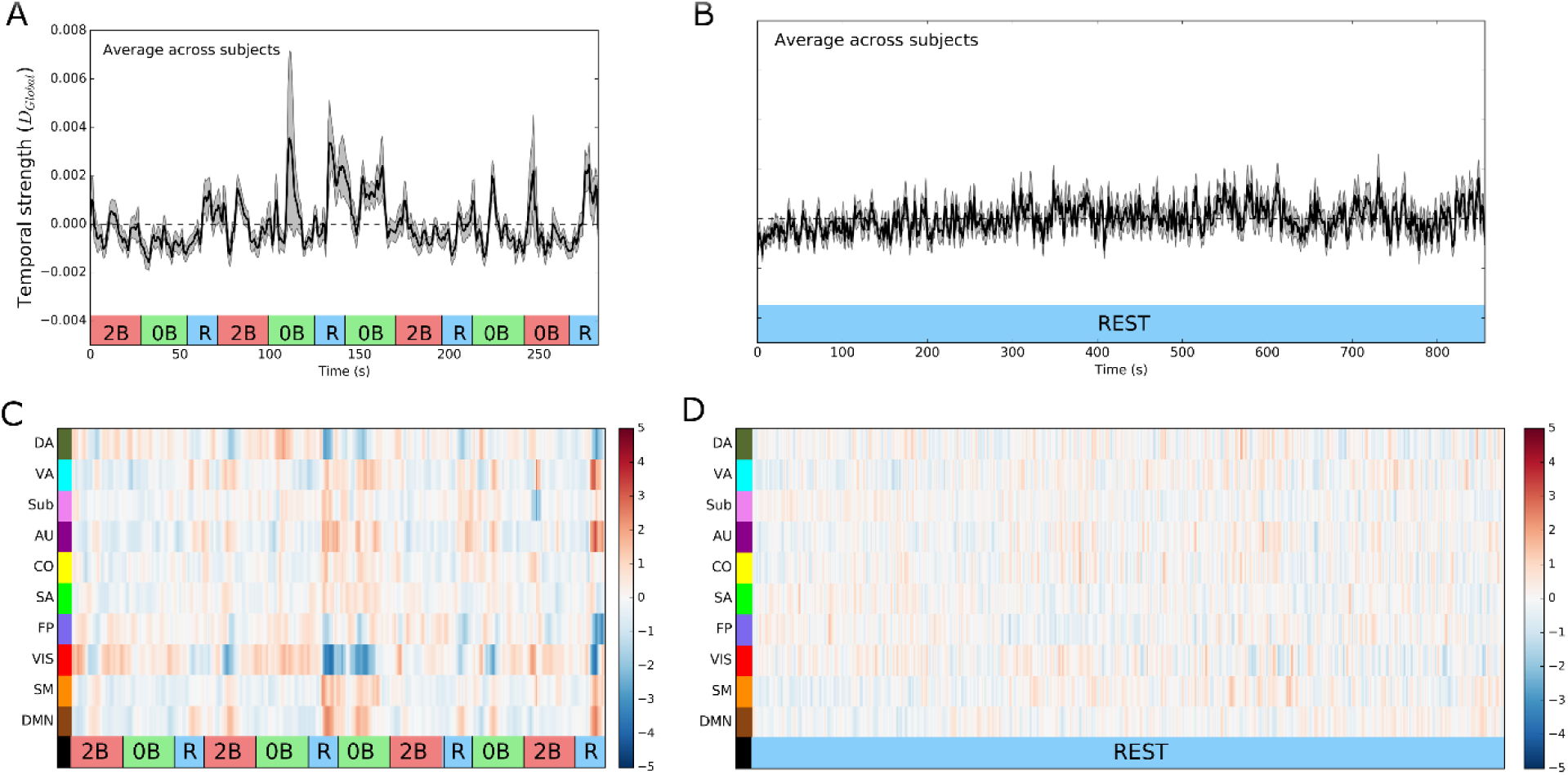
Replicability of the temporal strength results during the working memory and resting-state experiments. The results shown here compliments the results shown in Figure 2 and were computed on separate fMRI data with respect to the data used in main paper (the Right-Left (RL) dataset). (A,B) Global temporal strength (D_global_) averaged across all subjects during the working memory (A) and resting-state (B) experiments. (C,D) Temporal strength computed at a subnetwork level for the working memory (C) and resting-state experiments (D). ‘0B’ – 0-back epoch, ‘2B’ – 2-back epoch, ‘R’ – rest (baseline) epoch.

**Fig S3.**
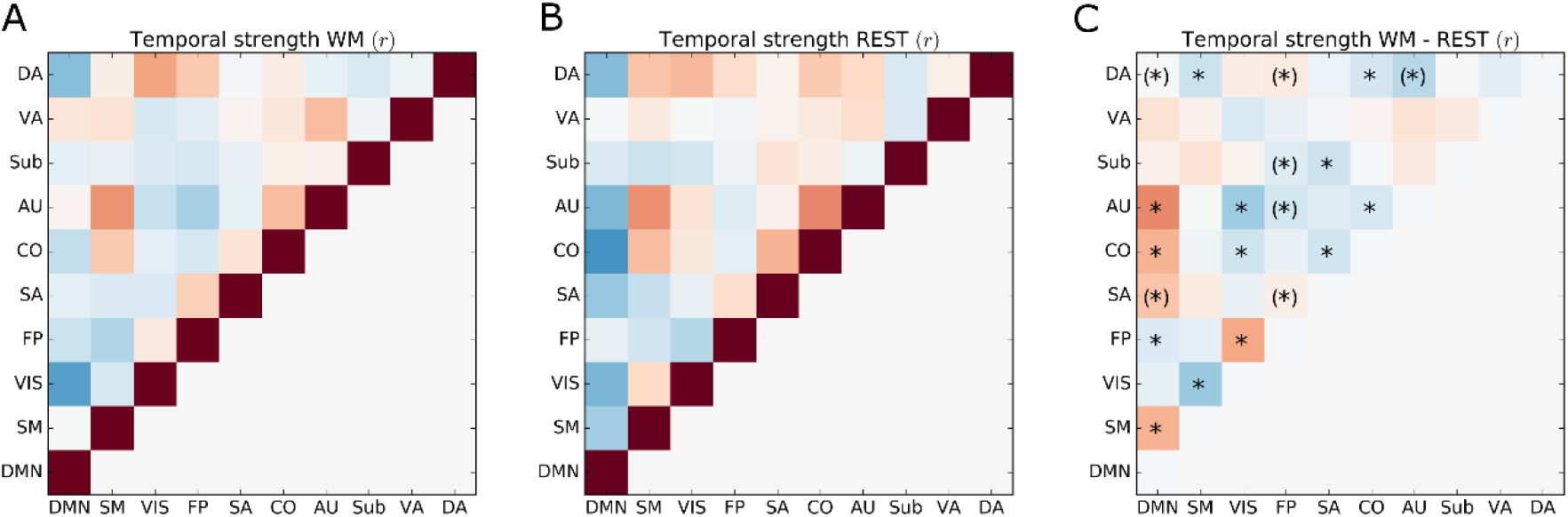
(A,B) Replicability of the correlation of temporal centrality time-courses between subnetworks during the working memory experiment (A) and resting-state (B) for the ‘RL’ data sets. (C) Difference in correlation of temporal strength for the working memory and resting-state experiments. Statistical significant differences in correlation were assessed by a non-parametric permutation approach (10000 permutations, p<0.001, Bonferroni corrected). This figure compliments the results shown in Figure 3. An asterisk within parenthesis signify that the differences in temporal strength time-courses where only significant in one of the datasets (‘LR’ and ‘RL’, respectively).

**Fig. S4.**
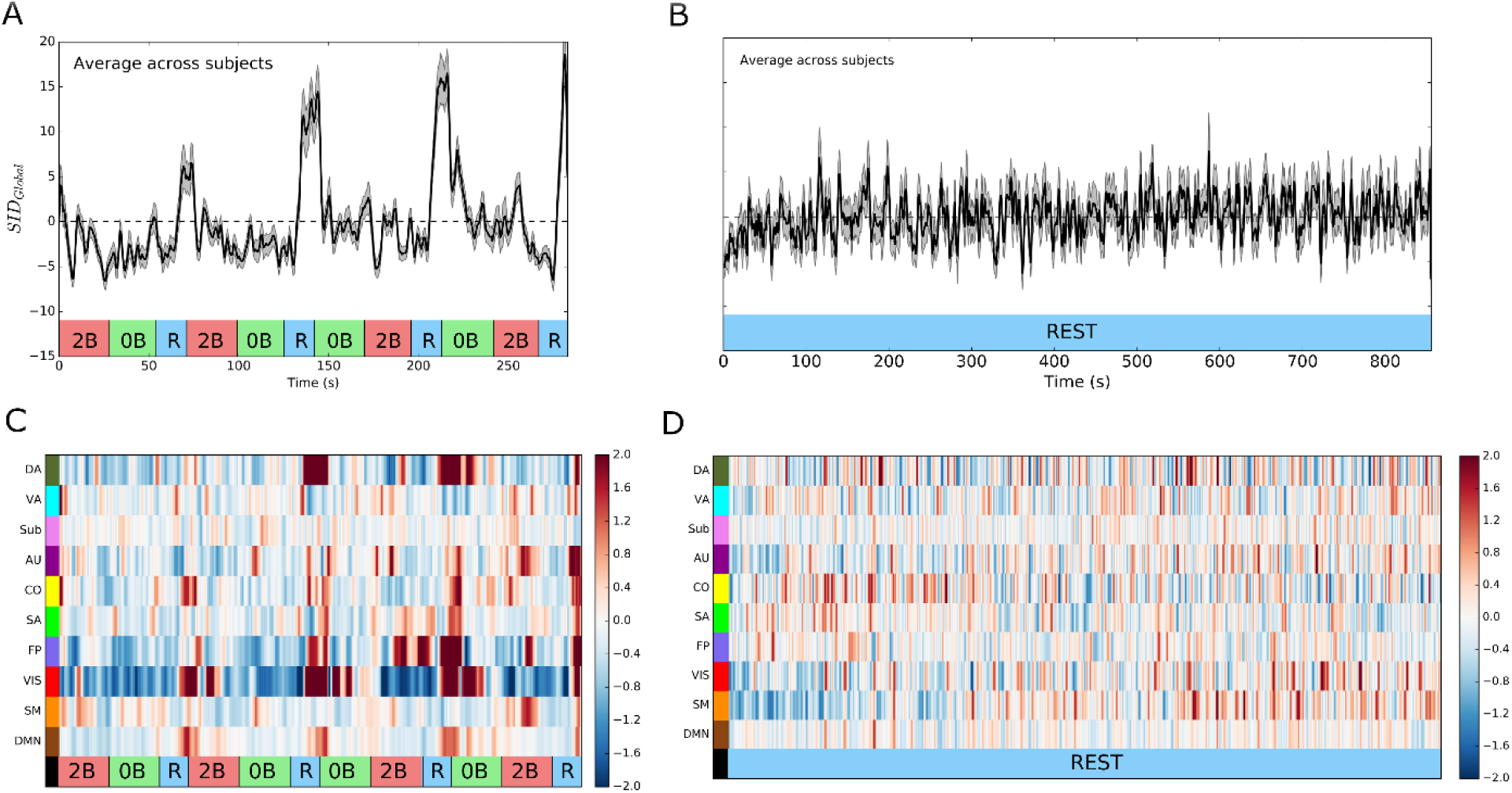
Replicability study of the Segregation Integration Difference (SID) results for the ‘RL’ fMRI datasets. (A,B) SID values at a global level (averaged across all subnetworks and subjects) during the working memory (A) and resting-state experiments (B). (C,D) SID at a subnetwork level (averaged across subjects) during the working memory (C) and resting-state experiments (D). ‘0B’ – 0-back epoch, ‘2B’ – 2-back epoch, ‘R’ – rest (baseline) epoch. Shaded areas depict the standard error of the mean. This figure compliments the results shown in Figure 4.

**Fig. S5.**
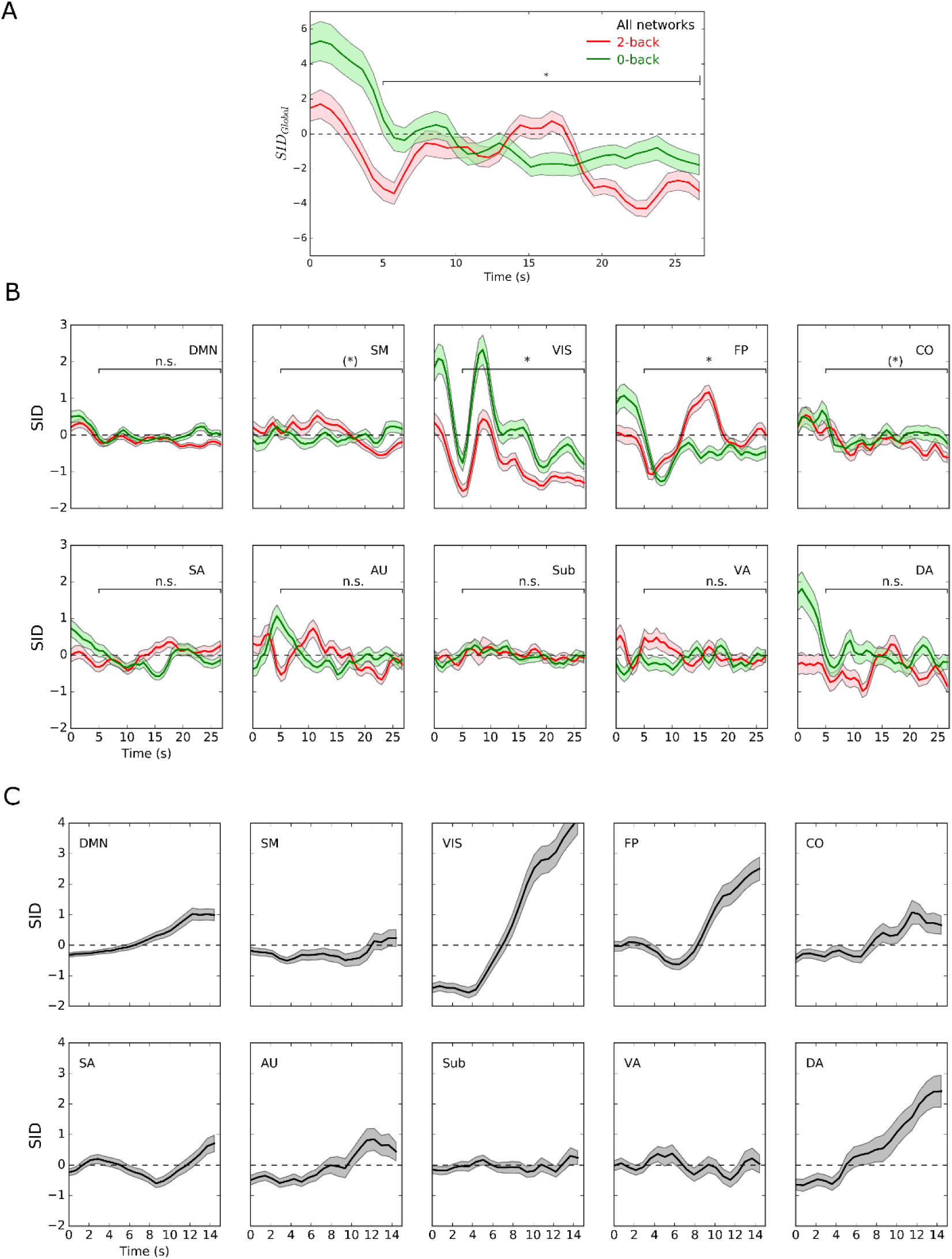
Replicability study of the time-locked average of SID results during 2-back, 0-back and baseline epochs for the ‘RL’ fMRI data sets. (A) Average of the global Segregation Integration Difference (SIDGlobal) values taken across all 2-back and 0-back epochs in all subjects for the working memory experiment. (B) Similar comparison as in (A) but here SID values were computed at the level of individual subnetworks. (C) SID plotted for the 15 seconds long baseline epochs averaged across baseline epochs and subjects. An asterisk within parenthesis signify that the differences in SID time-courses where only significant in one of the datasets (‘LR’ and ‘RL’, respectively). Shaded areas depict the standard error of the mean. The results shown in this Figure compliments the findings displayed in Figure 5.

**Fig. S6.**
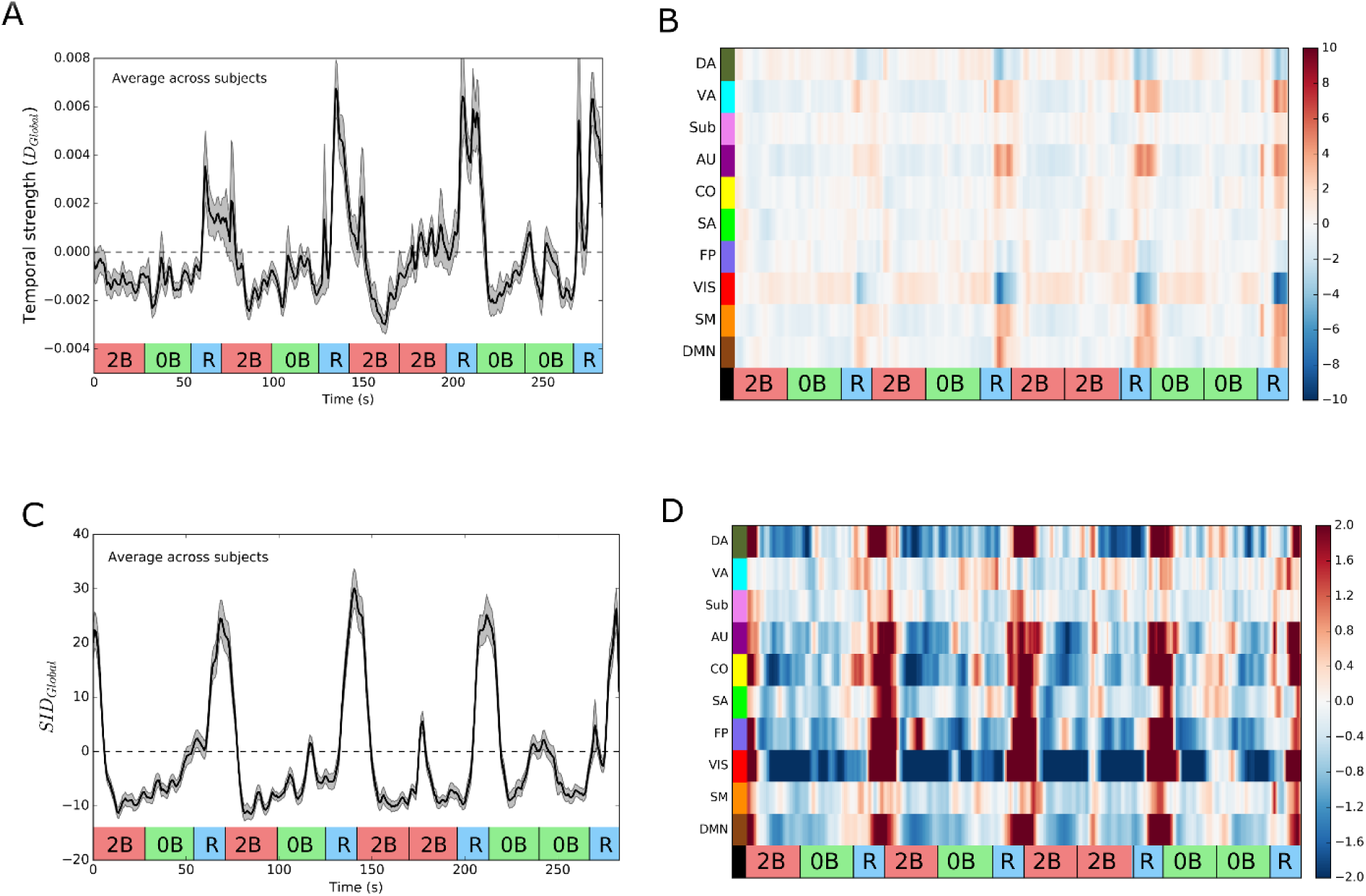
Temporal strength and SID during the working memory and resting-state experiments in the case when the jackknife correlation (JC) method to estimate time-varying functional connectivity was replaced with the spatial distance method (SD, described in detail in ref. 24). Apart from the change of method to compute time-varying functional connectivity time-series, calculations to derive estimates of temporal strength and SID were kept identical.). ‘0B’ – 0-back epoch, ‘2B’ – 2-back epoch, ‘R’ – rest (baseline) epoch. Shaded areas depict the standard error of the mean. The results shown in this Figure supplements the findings shown in Figures 2 and 4 and Supplementary Figures S2 and S4.

**Fig. S7.**
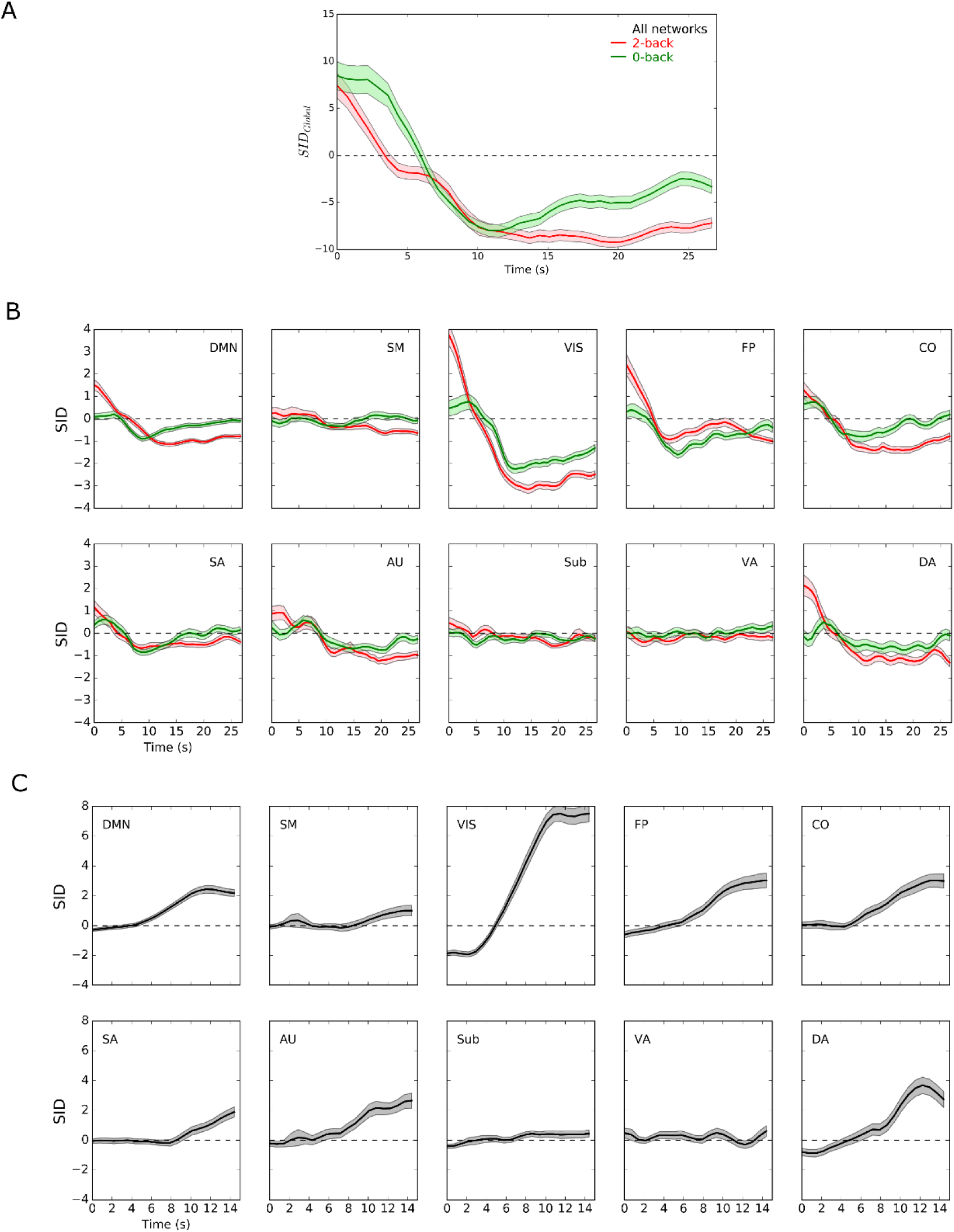
Results obtained when the jackknife correlation method was replaced with the spatial distance method (SD). (A) Time-locked averaging of 2-back, 0-back and baseline epochs taken across all ten subnetworks or for individual subnetworks (B-C). (A) Average of the SIDGlobal measure taken across all 2-back and 0-back epochs for all subjects in the working memory experiment. The results in this figure compliments the results shown in Figure 5 and Supplementary Figure S5.

**Fig. S8.**
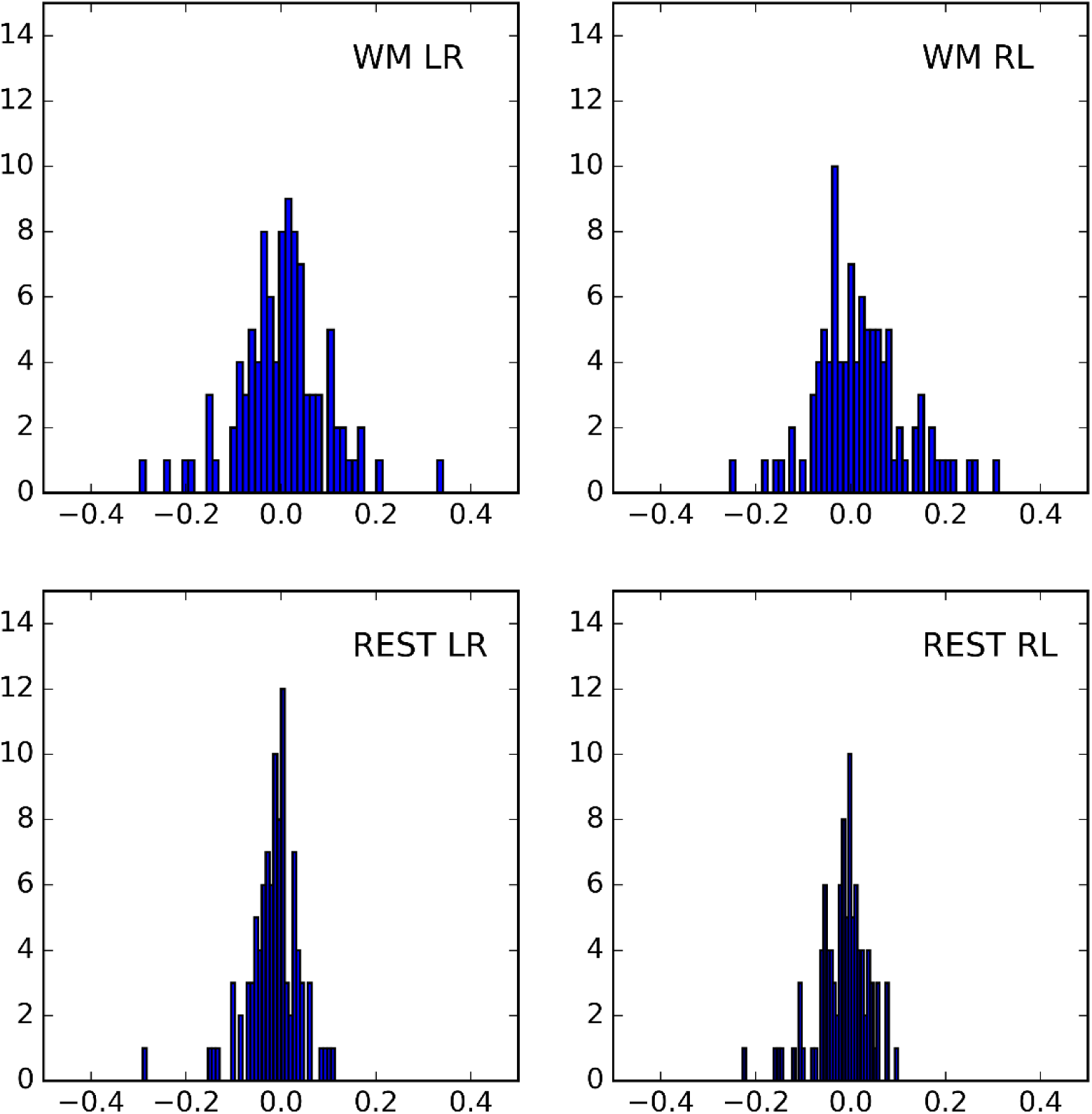
Distribution of correlation values for correlation between estimates of global temporal strength (D_global_, average over all nodes) and framewise displacement for all four fMRI sessions, (A) WM ‘LR’ data set, (B) WM ‘RL’ data set, (C) resting-state ‘LR’ data set and (D) resting-state ‘RL’ data set.

**Table S1.** DIC (Deviance Information Criterion) and ΔDIC values for three different drift diffusion models of the 2-back trials in the working memory experiment were fitted with respect to the SID values (i.e. whether the drift rate, decision boundary, or both relate to SID). Two different model configurations were used. The first included SID_Global_ estimates as a covariate that could change with different drift diffusion parameters. The second model configuration had SID_VIS_ and SID^FP^ estimates that could co-vary.

| Global SID model | Specification | DIC | delta DIC |
| --- | --- | --- | --- |
|  | v,a | 9698,0 | - |
|  | v | 9707,6 | 9,6 |
|  | a | 9837,1 | 140,1 |
| VIS+FP SID model | Specification | DIC | delta DIC |
|  | v | 9698,7 | - |
|  | v,a | 9702,0 | 3,3 |
|  | a | 9843,3 | 144,6 |

